# ModelMatcher: A scientist-centric online platform to facilitate collaborations between stakeholders of rare and undiagnosed disease research

**DOI:** 10.1101/2021.09.30.462504

**Authors:** J. Michael Harnish, Lucian Li, Sanja Rogic, Guillaume Poirier-Morency, Seon-Young Kim, Undiagnosed Diseases Network, Kym M. Boycott, Michael F. Wangler, Hugo J. Bellen, Philip Hieter, Paul Pavlidis, Zhandong Liu, Shinya Yamamoto

## Abstract

Next-generation sequencing is a prevalent diagnostic tool for undiagnosed diseases, and has played a significant role in rare disease gene discovery. While this technology resolves some cases, others are given a list of possibly damaging genetic variants necessitating functional studies. Productive collaborations between scientists, clinicians, and patients can help resolve such medical mysteries, and provide insights into *in vivo* function of human genes. Furthermore, facilitating interactions between scientists and research funders, including non-profit organizations or commercial entities, can dramatically reduce the time to translate discoveries from bench to bedside. Several systems designed to connect clinicians and researchers with a shared gene of interest have been successful. However, these platforms exclude some stakeholders based on their role or geography. Here we describe ModelMatcher, a global online matchmaking tool designed to facilitate cross-disciplinary collaborations, especially between scientists and other stakeholders of rare and undiagnosed disease research. ModelMatcher is integrated into the Rare Diseases Models and Mechanisms Network and Matchmaker Exchange, allowing users to identify potential collaborators in other registries. This living database decreases the time from when a scientist or clinician is making discoveries regarding their genes of interest, to when they identify collaborators and sponsors to facilitate translational and therapeutic research.

## Introduction

Rare and undiagnosed diseases are prevalent and costly issues facing the modern healthcare landscape. In the United States alone, 30 million individuals are estimated to be suffering from a rare disease, over 80% of which are estimated to have strong genetic components (Nguengang Wakap et al. 2020). The use of whole-exome sequencing (WES) or whole-genome sequencing (WGS) technologies as diagnostic tools has been increasing steadily. This increase is in part due to the decreasing cost of high-throughput sequencing. What remains a challenge is that many patients who undergo sequencing are still not given a specific diagnosis (Coventry et al., 2010; Lupski et al., 2011). Patients with an inconclusive WES or WGS result typically receive a list of several identified rare variants that are potentially responsible for the patient’s disease. While some variants can be ruled out by comparing the disease-gene association to the patient’s clinical presentation, this method still often leaves many candidates. A number of these genetic changes are variants of unknown significance (VUS) in known disease-causing genes whose associate phenotype has some overlap with the patient’s clinical presentation. Some are rare variants in genes of uncertain significance (GUS), which have yet to be associated with a genetic disorder in human. Missense variants in particular are difficult to interpret as modern *in silico* tools are still not able to accurately predict their functional consequences (Ghosh et al. 2017). To resolve these cases, experiments must be performed in a laboratory setting to better understand the functional consequence of the variant allele (Harnish et al. 2019). However, most clinicians and patients do not have contact with scientists who can carry out this task.

Collaborations between scientists and clinicians are increasingly commonplace, especially between model organism biologists and physicians that see undiagnosed patients (Wangler et al., 2017). Several countries and regions have created research programs designed to facilitate these endeavors. These programs include the Undiagnosed Diseases Network (UDN, https://undiagnosed.hms.harvard.edu/) in the United States (Gahl et al., 2016) and the Rare Diseases Models and Mechanisms (RDMM, http://www.rare-diseases-catalyst-network.ca/) Network in Canada (Boycott et al., 2020). Over the last five years, these platforms have been demonstrated to be very effective in facilitating diagnosis as well as identifying potential therapeutic avenues for many rare diseases. Functional studies in the UDN are carried out through the centralized Model Organisms Screening Centers (MOSC) by a relatively small number of model organism experts with broad biological knowledge. The MOSC sites utilize sophisticated genetic techniques in fruit fly, nematode worm, and zebrafish to characterize the effects of variant alleles found in rare disease patients to ultimately aid in the diagnosis and obtain mechanistic insights to further understand disease pathogenesis (Baldridge et al., 2021). While this UDN-MOSC system has many advantages including economy of scale, pipeline standardization, and establishment of strong trust relationships between scientists and clinicians over time, it doesn’t take full advantage of the entirety of scientific expertise that various scientists in the field possess. In contrast, RDMM adopts a decentralized model in which a Coordination Center that is supported by Clinical and Scientific Advisory Committees performs matchmaking between model organism scientists and clinicians with shared genetic interests within Canada (Boycott et al., 2020). Once a match is made, scientists are given seed grants (Can$25,000/case) to carry out experiments. Over the past three years, similar platforms were established in Europe [Solve-RD (https://solve-rd.eu/)], Japan [J-RDMM (https://j-rdmm.org/indexEn.html)] and Australia [Australian Functional Genomics Network (AFGN, https://www.functionalgenomics.org.au/) to facilitate collaborative research within each county or region. The RDMM model catalyzes and supports collaborations within each network, and there has been some effort to facilitate collaborations across different RDMM-like networks. However, there are many scientists and clinicians who cannot participate (e.g., in countries like the United States), primarily because there is a large administrative hurdle in transferring grant funds across borders. Since clinicians with undiagnosed patients and scientists with expertise in specific genes may be found in different parts of the world, there is an unmet need to establish an open platform designed to facilitate collaborations on a global scale. In addition, while these existing platforms have been successful in facilitating collaborations between model organism biologists and clinical groups, scientists with other types of expertise such as cell biologists, biochemists, and structural biologists who primarily carry out *in vitro* experiments are under-represented.

In addition to clinicians, other stakeholders of rare and undiagnosed diseases such as patients, families, non-profit entities, public funding agencies, and biotech or pharmaceutical companies can benefit from having direct connections with scientists. In contrast to clinicians who often interact with these entities, only a handful of scientists, most of whom are affiliated with medical schools, have experience in interacting with these types of stakeholders. Considering that there are many scientists in basic biology departments who have the capacity to conduct sophisticated gene and variant functional studies in a timely manner, it will be valuable to establish a network in which both clinical and non-clinical stakeholders can match with scientists who possess relevant expertise. Such a network will enable all stakeholders to find scientists with expertise in specific genes and disciplines to collaborate with them in order to attempt to diagnose, understand, and ultimately manage the rare disease of interest. Scientists also stand to benefit from such a system because collaborations demonstrating the relevance of their research to human disease can often increase the significance and impact of their ongoing research projects, which can in turn help secure additional sources of funding for their laboratories. Furthermore, data obtained from studying rare variants identified in patients can also provide scientists with new information about their genes of interest, expanding fundamental knowledge that has a broad impact beyond the specific patient and disease.

Finally, interdisciplinary collaborations between scientists are becoming more and more critical to make significant advances in the field of biomedical sciences. In contrast to several decades ago when single laboratory papers were commonly published in high-impact journals, collaborative manuscripts with contributions from multiple laboratories with a wide range of expertise now dominate these journals (Sekara et al., 2018). Collaborations between scientists are often established locally, or through personal networks that are built based on face-to-face meeting opportunities including scientific conferences. While social media and virtual scientific networks such as Google Scholar (https://scholar.google.com/) and ResearchGate (https://www.researchgate.net/) can also be utilized to identify potential collaborators, the vast amount of information on these sites makes identifying the right collaborator a daunting task. Facilitating collaborations between different types of scientists with interests in the same or orthologous genes in multiple species has been referred to as ‘vertical integration’ (https://orip.nih.gov/sites/default/files/Validation_Session_IX_Meeting_Report_Final_508.pdf). As an example of vertical integration, studies of orthologous genes in fruit flies, zebrafish, and human have provided complementary information to provide a diagnosis for rare disease patients (Barish et al., 2020; Ravenscroft et al., 2021). Furthermore, multidisciplinary studies by model organism biologists, cell biologists and biochemists have successfully classified the pathogenicity of variants in conserved evolutionarily genes in the context of both rare and common diseases (Post et al., 2020). While open services that include scientist-clinician matchmaking as part of their larger scope already exist such as GeneMatcher (https://genematcher.org/) (Sobreira et al. 2015), scientist-scientist matchmaking is rarely performed on such platforms.

After considering the above points carefully and discussing with many stakeholders, we concluded that a new platform was needed to address multiple gaps in the current biomedical collaboration ecosystem. To this end, we developed ModelMatcher (https://www.modelmatcher.net/) to address the unmet need of an open, international, decentralized network devoted to pursuing functional studies relevant to rare and undiagnosed diseases which is catered to all stakeholders (**Table 1**). The core of ModelMatcher is an open registry of scientists who input their genes of interest as well as their experimental and organ system expertise. Most information on this database is publicly searchable, but we also offer a matchmaking system that allows registered users to reach out to scientists with unpublished or confidential information. ModelMatcher is further connected to external scientist and clinical registries through the RDMM International and Matchmaker Exchange (MME) networks, respectively. These developments will expand the network paradigm for biomedical research by extending the collaboration ecosystem worldwide and including rare disease stakeholders, such as patients and funding groups, who are currently un- or under-represented in the current collaborative ecosystem.

**Table 1.** Features and strengths of networks and tools that facilitates collaboration between rare and undiagnosed disease stakeholders. Comparison of key features and strengths of the Undiagnosed Disease Network (UDN), Rare Disease Models and Mechanism (RDMM) Network, Matchmaker Exchange (MME), PubMed (literature search), and ModelMatcher are listed here.

| Collaboration platform | Decentralized structure | Up to date information | No geographic barrier | Participating rare disease stakeholders |  |  |  |
| --- | --- | --- | --- | --- | --- | --- | --- |
|  |  |  |  | Clinicians | Scientists | Patients | Research funders |
| UDN |  | X |  | X | X | X | X |
| RDMM | X | X |  | X | X |  | X |
| MME | X | X | X | X | X | X |  |
| PubMed | X |  | X | X | X |  |  |
| ModelMatcher | X | X | X | X | X | X | X |

## Results

### The scientist registry serves as the starting point for all collaborations through ModelMatcher

The primary function of ModelMatcher is to provide users with a list of collaborative scientists who have expertise in specific human genes or their ortholog(s) in genetic model organisms (**Figure 1**). ModelMatcher accomplishes this by maintaining an active Scientist Registry based on a software that was originally developed for the RDMM (Boycott et al., 2020). The information stored in this registry includes the scientist’s name and contact information, affiliation, principal investigator (PI) status, research interests, relevant publications, types of experiments that are performed in their laboratories (e.g., *in vitro, in vivo, in silico*, etc.), organisms they use to conduct gene function studies, organ systems they have expertise in, and their genes of interest (**Figure 2A**). While the original RDMM software was primarily designed to recruit model organism scientists who conduct *in vivo* experiments, other types of scientists including cell biologists, molecular biologists, biochemists, structural biologists, and computational biologists are also encouraged to participate in ModelMatcher (**Figure 2B, B’**). If a scientist works with one of the ten supported model organisms [human (*Homo sapiens*), mouse (*Mus musculus*), rat (*Rattus norvegicus*), frog (*Xenopus tropicalis*), zebrafish (*Danio rerio*), fruit fly (*Drosophila melanogaster*), nematode worm (*Caenorhabditis elegans*), budding yeast (*Saccharomyces cerevisiae*), fission yeast (*Schizosaccharomyces pombe*), or *E. coli* (*Escherichia coli*)], they can enter their genes of interest based on official gene symbols for each species (**Figure 2C-D’**). If they work with other species, scientists are encouraged to provide the gene symbol of the orthologous human gene, which could be identified based on information from databases such as Ensembl (https://useast.ensembl.org/) or homology alignment programs such as BLAST (https://blast.ncbi.nlm.nih.gov/). If a scientist works on multiple model organisms, this information is also captured in the Scientist Registry.

**Figure 1.**
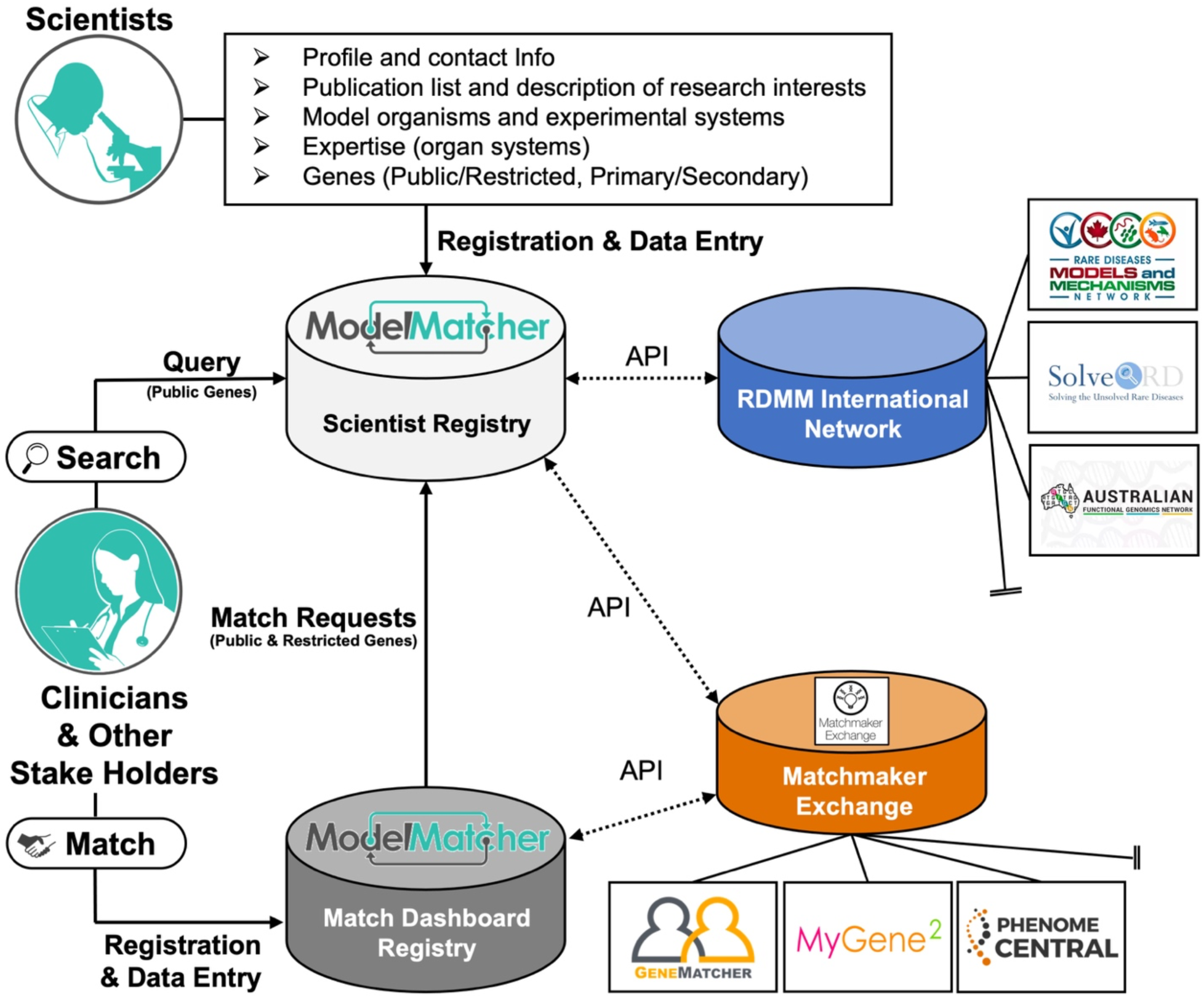
Basic structure of ModelMatcher and its relationship to partner registries. Scientists who wish to identify potential collaborators and sponsors register their information into the Scientist Registry of ModelMatcher. In addition to basic information about their interests and expertise, scientists enter their genes of interest, which are classified into Public or Restricted privacy categories and Primary or Secondary priority tiers (see text for additional information about gene classification). Clinicians and other stakeholders of rare and undiagnosed disease research can mine this data in two ways. The Search feature allows anyone to query the information associated with genes classified as Public without needing to register in ModelMatcher. In addition to identifying scientists in the ModelMatcher Scientist Registry, Search feature users will also receive information about scientists in partner scientist registries that are part of the RDMM International Network including RDMM, Solve-RD, and AFGN. The Match Request feature, which requires users to register for a Match Dashboard account, provides a unique mechanism for users to contact a scientist who not only classified a gene as Public but also Restricted. The Match Dashboard account also enables users to track their matches, provides a communication platform to send messages to scientists, and further provides an entry point to connect with clinicians and patients who participate in clinical registries that are part of Matchmaker Exchange including GeneMatcher, MyGene2 and PhenomeCentral.

**Figure 2.**
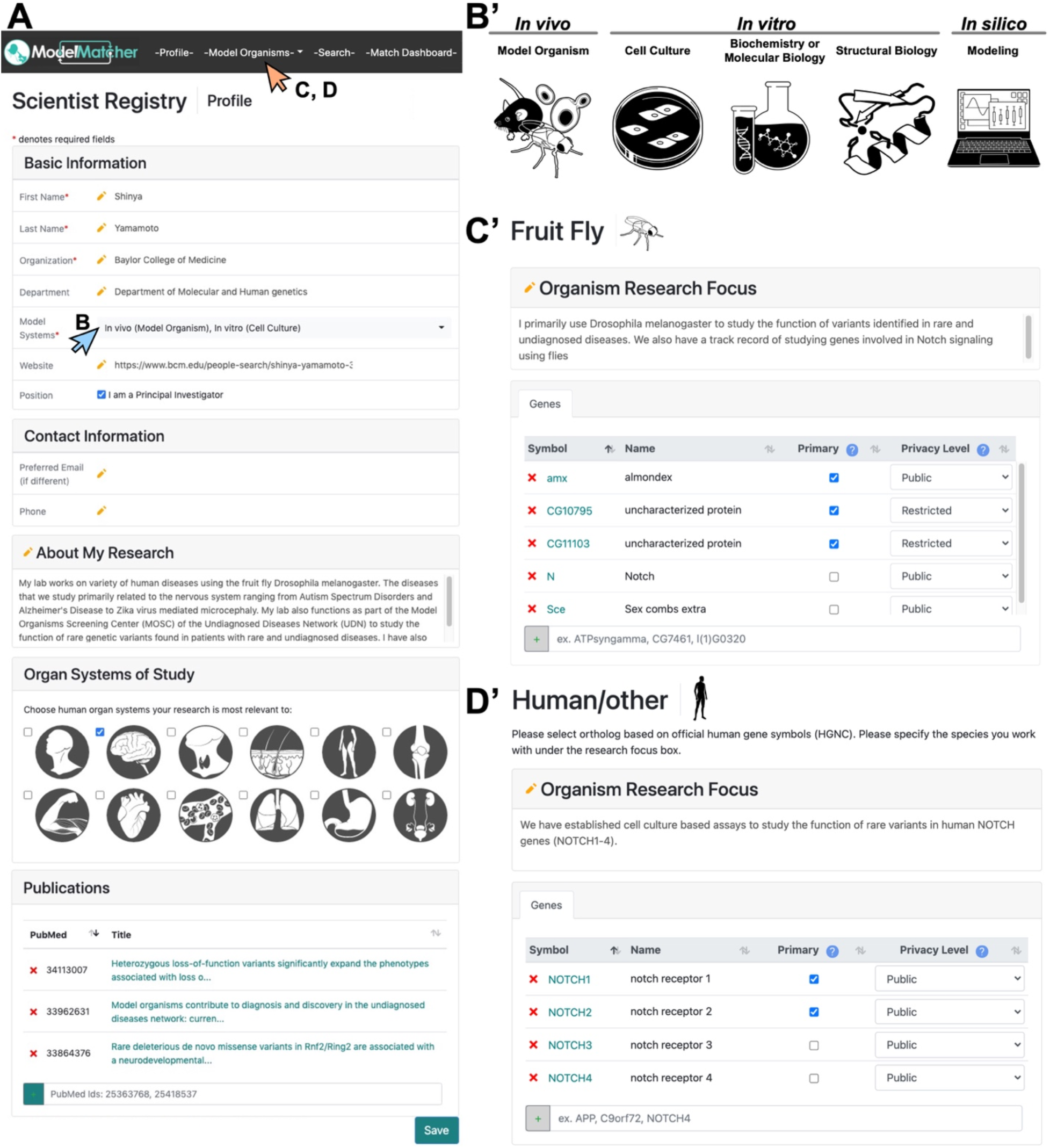
Information collected from scientists to establish the ModelMatcher Scientist Registry. Upon registration, ModelMatcher requires scientists to enter information about themselves and the genes they are interested in. **A)** The -Profile- tab requires scientists to provide general information, including who they are, where they work, whether they are PIs of their labs, and how to contact them. It also collects information about their research interest, what kind of experiments they conduct (see **B**), what organ systems they have expertise in, and any relevant papers they have published. **B-B’)** Upon clicking the “Model Systems” pull down menu in **A)**, scientists can select one or more experimental model systems that they use to perform functional studies of genes and genetic variants. **C-D)** By clicking the “Model Organisms” pull down menu in **A)**, scientists can select the model organism they work on to access the gene entry pages. **C’)** shows a gene entry page for fruit fly (*Drosophila melanogaster*) and **D’)** shows a gene entry page for human or other species that are not currently supported by ModelMatcher. Here, scientists can enter information about their specific research topic using each organism and generate a list of genes they are willing to collaborate on using the official gene symbol that is specific to each species. Scientists can categorize each gene into Primary or Secondary priority tiers based on whether they are actively working on the specific gene or not. Furthermore, scientists can classify genes into Public or Restricted privacy categories based on confidentiality levels. Public genes will be associated with the scientist in both the Search and the Match Dashboard features. Restricted genes will show up as a gene that is entered by an Anonymous Scientists, but will not divulge the information of the individual who registered it. The identity of the Anonymous Scientist will only be revealed if the scientist wishes to disclose the information upon receiving a Match Request email from individual Match Dashboard or Matchmaker Exchange users. This allows scientists to identify potential collaborators on unpublished projects that consider as confidential due to potential competitors in their field.

While most scientists are open to publicly declaring their genes of interest, some scientists with unpublished data may only wish to share such information with potential collaborators to avoid being scooped by their competitors. Therefore, we integrated a gene level privacy feature in which a scientist can classify their genes of interest into Public and Restricted categories on a case-by-case basis. Public genes will be searchable by anyone that has access to the open ModelMatcher website, whereas scientists themselves can control access to information regarding their Restricted genes, which will be discussed in more detail in the following section. In addition, scientists are also asked to classify their genes of interest into Primary and Secondary priority tiers. Primary tier genes are genes that a scientist’s lab actively works on, and has protocols and reagents ready to perform functional assays. In contrast, Secondary tier genes are genes that a scientist’s lab has worked on in the past or has a strong interest in, but would need additional time to establish protocols or acquire/develop reagents to conduct experiments. Such information can be used to estimate how quickly a scientist may be able to generate data, which could become important when a quick turnaround of functional data is required.

### The Search function allows anyone to query public data from ModelMatcher and partner scientist registries

The Scientist Registry of ModelMatcher can be queried by anyone using the Search feature that does not require user registration (https://www.modelmatcher.net:8443/search, **Figure 3**). Users of this feature can input a gene of interest and optionally narrow their results via placing filters on model organism, organ system, or gene tier if desired (**Figure 3A**). The default setting prompts the users to input a human gene of interest based on HGNC (HUGO Gene Nomenclature Committee, https://www.genenames.org/) nomenclature (Tweedie et al., 2021) and returns a list of scientists with expertise in this human gene or their ortholog candidates across all supported model organisms. Ortholog candidates of human genes are primarily identified based on the DIOPT (DRSC Integrative Ortholog Prediction Tool, https://www.flyrnai.org/cgi-bin/DRSC_orthologs.pl), an integrative ortholog prediction tool that uses 18 different algorithms and provides a score that reflects the likelihood of certain gene pairs to be orthologous to one another (Hu et al., 2011) (**Figure 3B**). Some gene pairs that are true orthologs with low DIOPT scores have been manually included, in addition to a list of orthologs for model organisms that are not currently supported by DIOPT [e.g., *E coli*]. As improved lists of true orthologs become available through community efforts, such as the Alliance of Genome Resources (Agapite et al., 2020) and Quest for Orthologs (Altenhoff et al., 2021), we plan to adopt such datasets to improve the accuracy of ortholog identification in order to remove false-positive results while including false-negative data points. Basic information for each scientist identified as having expertise in the specific human gene or their ortholog candidates will be displayed in a simple search output table (**Figure 3C**). Users can further navigate to the scientist’s profile page to find additional information about their research in order to identify the ideal collaborator candidate. Contact information is listed under each user’s profile page, which can be used to initiate a conversation via email. Some scientists may be listed more than once if they work on orthologous genes in multiple species. When there is a scientist who has classified the gene of interest in their Restricted category, such information will be displayed as information from an Anonymous Scientist. In order to reach out to an Anonymous Scientist, users will need to create a Match Dashboard account, which is discussed below.

**Figure 3.**
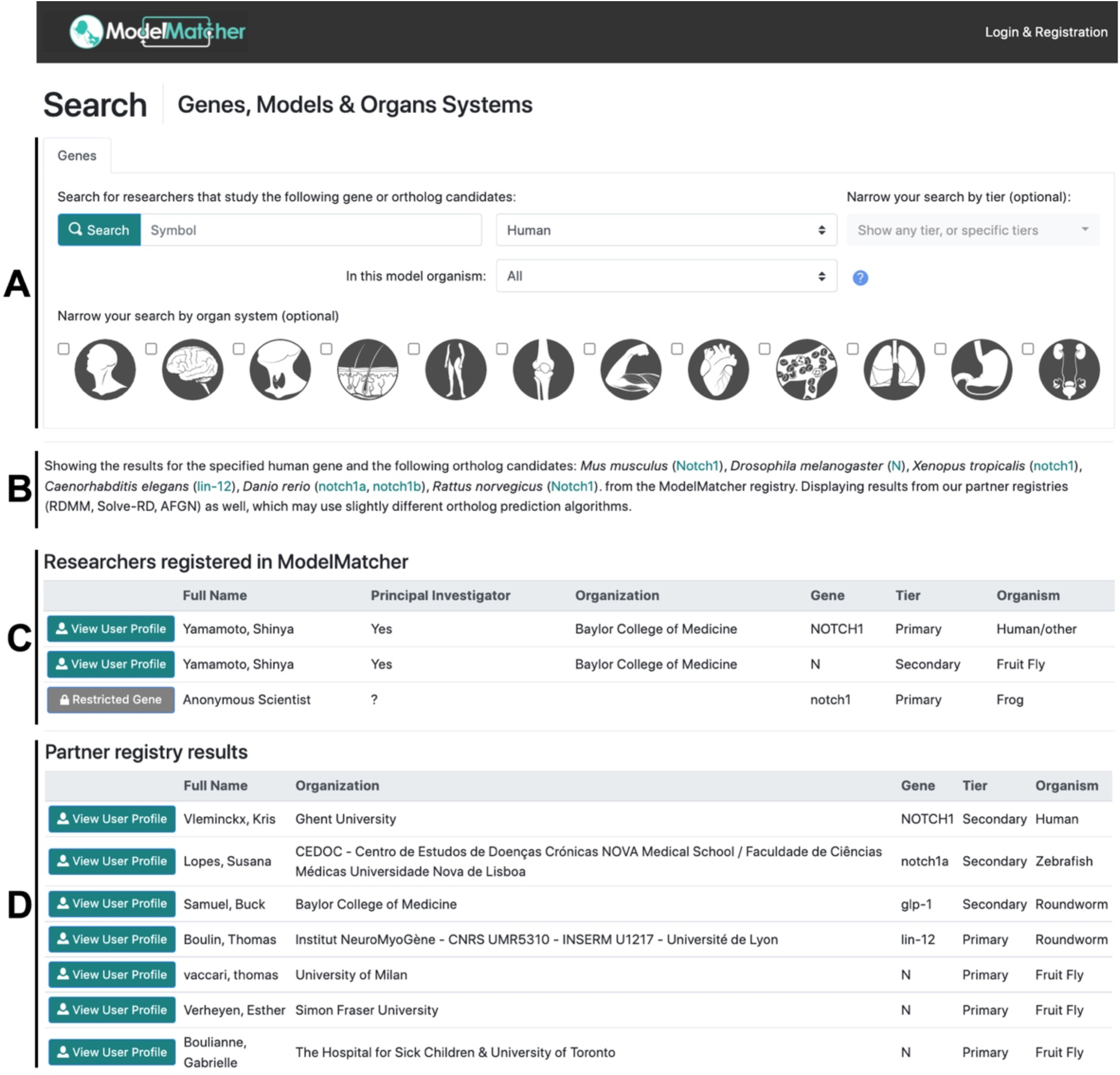
Search feature identifies scientists registered in ModelMatcher and partner registries. The Search page outputs information on scientists who work on specific human genes or its predicted orthologs in model organisms. **A)** A query is typically initiated by entering a human gene symbol according to the official nomenclature, although one can initiate a search based on model organism genes if desired. If necessary, searches can be refined by which model organism is being used, whether the gene is classified as Primary or Secondary priority tiers, and in which organ system the scientist specializes. **B)** Upon clicking the “Search” button in A, a list of genes considered ortholog candidates of the human gene of interest (human *NOTCH1*, in this case) in the ten supported organisms is be displayed. **C)** When a Search is performed, a table of ModelMatcher scientists who registered the human gene of interest or its ortholog candidates is generated. If the gene is classified as Public, the name of the scientist, their affiliation, and whether they are PIs or not will be displayed along with the matched gene, gene’s priority tier and the corresponding model organism. Users can click the “View User Profile” button to access their full user profile page (see **Figure 5B**). A scientist who classified the specific gene as Restricted will be shown as an Anonymous Scientist, who can be contacted through the Match Dashboard (see **Figures 4-5**). **D)** When a Search is performed, a table of scientists registered in partner registries in the RDMM International Network (RDMM, Solve-RD and AFGN) who expressed interest in the human gene of interest or its ortholog candidates is also generated.

In addition to querying the Scientist Registry of ModelMatcher, our Search feature allows users to query the list of scientists within the RDMM International Network (**Figure 1**). This network consists of multiple scientist registries which have been built using the same RDMM software. These registries communicate with each other via APIs (application programming interfaces) in real time when a query is made. Currently, information that is classified as Public from RDMM, Solve-RD, and AFGN is displayed below the table of ModelMatcher scientists in a similar manner (**Figure 3D**). By clicking the “View User Profile” button for each entry, users will be able to access information about scientists in partner registries and reach out to them through the contact information listed there. Therefore, in addition to identifying scientists who registered their information in ModelMatcher, users can identify potential collaborators in Canada, Europe, and Australia who are already part of a regional rare disease research network.

### Match Dashboard provides additional features to query the Scientist Registry and initiate communication

In addition to the Search feature that allows anyone to query the ModelMatcher and partner registries as discussed above (**Figure 3**), we built the Match Dashboard feature that provides additional benefits to registered users (**Figure 4-5**). A Match Dashboard account, which is independent of a Scientist Registry account, can be created by anyone who wishes to collaborate with scientists through a separate registration webpage (https://www.modelmatcher.net/login.html). Upon registration, users gain access to the Match Dashboard, which is a personalized webpage that facilitates communication with potential collaborators. In addition, the Match Dashboard can be used to identify potential clinical collaborators through MME, which is discussed in the following section.

**Figure 4.**
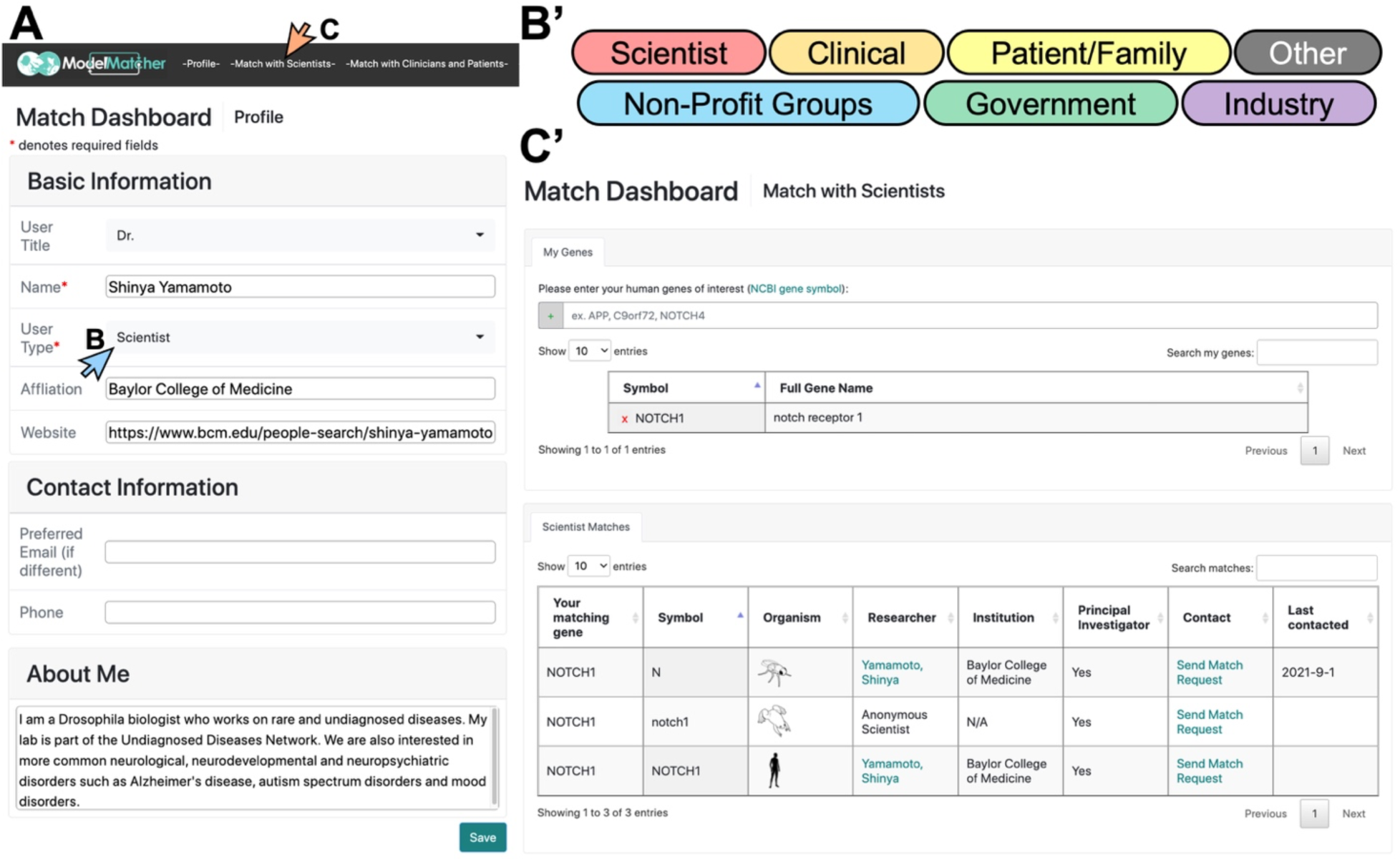
Information collected from rare and undiagnosed disease research stakeholders to populate a Match Dashboard account. Upon creating a Match Dashboard account, users enter information about themselves and the human genes they are interested in. **A)** The -Profile- tab requires users to provide general information, including who they are, where they work (if applicable), and why they are interest in rare and undiagnosed diseases. **B-B’)** Upon clicking the “User Type” pull down menu in **A)**, users can select one or more user categories to which they feel they belong. **C-C’)** The -Match with Scientists- button in **A)** will navigate the users to a tab to enter their human genes of interest to identify potential scientific collaborators. Upon entry, a table will be generated in the “Scientist Matches” box at the bottom of the -Match with Scientist- tab that lists all scientists who expressed interest in the same human gene or its ortholog candidates. Scientists who selected the specific gene as Public are identifiable whereas scientists who selected the gene as Restricted are displayed as an Anonymous Scientist.

In their profile page, Match Dashboard users input their information including their user category (e.g., patient/family member, clinician, scientist, industry, etc.), affiliation (if any), contact information, and a general statement regarding their interest in rare and undiagnosed diseases (**Figure 4A-B**). Once this is complete, the user will then navigate either to the -Match with Scientists- (**Figure 4C**) or -Match with Clinicians and Patients- tab (**Figures 4D, 6**). When the user clicks the -Match with Scientists- tab and enters the human genes they are interested in based on HGNC nomenclature, ModelMatcher generates a table of scientists who registered these human genes or their ortholog candidates from our Scientist Registry (**Figures 4C’, 5A**). This display in the Match Dashboard is similar to what is returned via the public Search function, although it provides additional features. First, the genes of interest list for each user is stored in our Match Dashboard registry, which is kept separately from the Scientist Registry, and queried against the latest Scientist Registry in real time. Therefore, whenever a user logs into their Match Dashboard account, they will be able to identify all of the matching scientists without having to perform individual searches. This is especially useful if a user has multiple genes of interest that they would like to keep an eye on longitudinally, considering that additional scientists with expertise in the gene of interest may appear in the future. Second, the user will have access to the Match Request Form, which is a messaging system that facilitates the communication process. Upon identification of a scientist whom the user wishes to contact based on the scientist’s profile (**Figure 5B**), the user can open the Match Request Form by clicking the “Send Match Request” button that is shown for each scientist (**Figure 5C**, arrow). A new tab will open that contains a contact form addressed towards the specific scientist regarding the specific gene (**Figure 5C’**). The user may add a specific reason for which they want to contact this scientist if this is not documented in the general user profile, and click the “Send” button to send a message. ModelMatcher will then generate a standardized email that will be sent to the scientist. The user’s profile from Match Dashboard is automatically included in this email, which saves the user from entering the same information every time they contact a new scientist. More importantly, Match Request Forms can also be used to contact Anonymous Scientists who classified specific genes as Restricted and whose contact information is not publicly displayed (**Figure 5D, 5D’**). When a message is sent to the scientist, the system logs the last contact date and displays this information on the Match Dashboard (**Figures 5A**), providing the user with an easy way to track which scientists they contacted, and how long ago that attempt to contact took place. In summary, the -Match with Scientists- feature of Match Dashboard allows users to reach out to scientists regardless of the gene privacy level using a standardized email template, and provides a tracking system to log all of the communication attempts.

**Figure 5.**
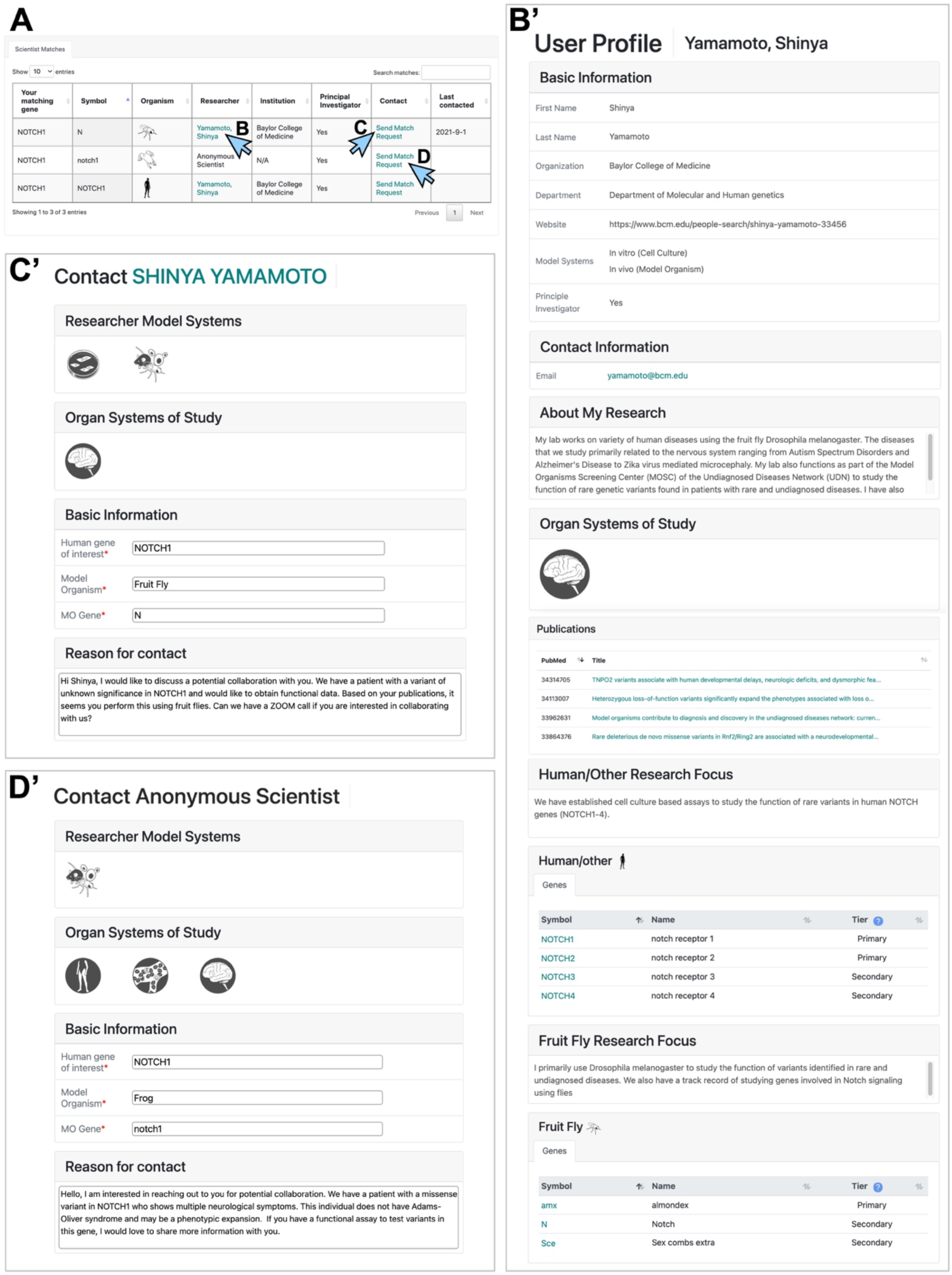
Match Dashboard allows users to contact scientists through a standardized Match Request Form. **A)** Starting from the table of scientists and their genes of interest shown in **Figure 4C’**, Match Dashboard users can obtain information about each scientist and utilize the Match Request Forms to initiate communication. **B-B’)** By clicking a scientist’s name shown teal color in **A)**, Match Dashboard users can access their full public profile page. Note that the same information can also be obtained from clicking the “View User Profile” button next to the corresponding scientist’s name in the Search result window shown in Figure 3C. **C-D)** By clicking the “Send Match Request” link in **A)**, a Match Request Form will be displayed that allow users to contact the specific scientist. **C’)** shows a contact form addressed to a scientist who categorized the gene of interest as Public, while **D’)** shows a contact form addressed to a scientist who categorized the gene as Restricted. Note that some non-identifiable information about the anonymous user, including whether they are a PI, their experimental model systems, and their organ systems of interest are displayed. The identity of the Anonymous Scientist will be revealed once the scientist responds to a specific user’s match request, and only to the person who made the match request.

### Integration of ModelMatcher into the Matchmaker Exchange network allows our users to discover clinical collaborators and vice versa

While the Search and Match Dashboard features allow ModelMatcher users to discover scientists with expertise in certain genes and experimental/organ systems, they do not allow our scientists to effectively discover clinical collaborators since they need to passively wait to be contacted by a clinician. In order to allow scientists to actively match with clinicians who have access to patients with rare, potentially pathogenic variants in their genes of interest, ModelMatcher became part of MME (https://www.matchmakerexchange.org/), a large network of federated databases for rare disease gene discovery (Philippakis et al., 2015). So far, ModelMatcher has developed partnerships with three nodes of MME: GeneMatcher, PhenomeCentral and MyGene2. GeneMatcher (https://genematcher.org/) is a matchmaking platform provided by the Baylor-Hopkins Center of Mendelian Genomics (CMG) in the United States (Sobreira et al. 2015). As of September 2021, more than 10,000 users, most of whom are clinicians or clinical researchers, from >90 countries have submitted >50,000 match requests corresponding to >13,000 human genes. PhenomeCentral (https://www.phenomecentral.org/) is a service developed in Canada that integrates both genotype and phenotype information to match clinicians with rare disease patients (Buske et al., 2015). As of September 2021, over 12,000 cases have been entered into this registry from research initiatives such as FORGE (Finding of Rare Disease Genes in Canada) (Beaulieu et al., 2014), Care4Rare (Care for Rare Canada) (http://care4rare.ca/), UDN (Brownstein et al., 2015) and UDN International (Taruscio et al., 2020). MyGene2 (https://mygene2.org/) is a patient-centric matchmaking platform hosted by the University of Washington CMG in the United States (Chong et al., 2016). As of September 2021, more than 4,000 families with patients with rare diseases have joined the network, many of whom are sharing their profiles with users from other MME nodes. Using standardized MME APIs that allow the connection of ModelMatcher to these and other collaboration focused network nodes of MME, users of ModelMatcher can identify clinicians and patients who share interest in the same or orthologous genes and vice versa.

ModelMatcher users can identify clinicians and patients from other MME nodes using their Match Dashboard account. By creating a Match Dashboard account as described above (**Figure 4A**), both scientists and non-scientists can take advantage of this feature. Once the user logs in to their Match Dashboard page, they can navigate to the -Match with Clinicians and Patients- tab to access this feature (**Figure 6**). From here, the user enters their human gene of interest and send Match Requests to other parties. The genes entered in the -Match with Scientists- tab will be automatically loaded into the -Match with Clinicians and Patients- so that the list of genes of interest will be synchronized within the Match Dashboard. When the user clicks the “Exchange Info” button shown for each gene, they will be taken to a Match Request Form that allows them to match with clinicians or other stakeholders interested in this gene from other MME nodes (**Figure 6B**). Users will be given the option to opt-out of matching with a specific registry if they feel that it is not applicable to their use case. When an MME match request is executed by clicking the “Send” button (**Figure 6C**), the user’s profile and additional information provided for the specific gene will be sent to the MME partner registries, which will then return information regarding their matching cases to ModelMatcher. We provide this match data to our users in two ways. First, users will receive an email detailing the case information of any external matches they have received. Even if the match results are negative, they will receive an email documenting this information, together with a recommendation to repeat the MME Match Request at a later time point (e.g., after ∼6 months). Second, a history of Match Requests for each gene is stored and displayed on the Match Dashboard (**Figure 6C’**), allowing users to access the information of previous matches without having to sort through their emails. This prevents the users from performing multiple redundant Match Requests in a short time frame, and allows them to keep track of previous matches for their genes of interest.

**Figure 6.**
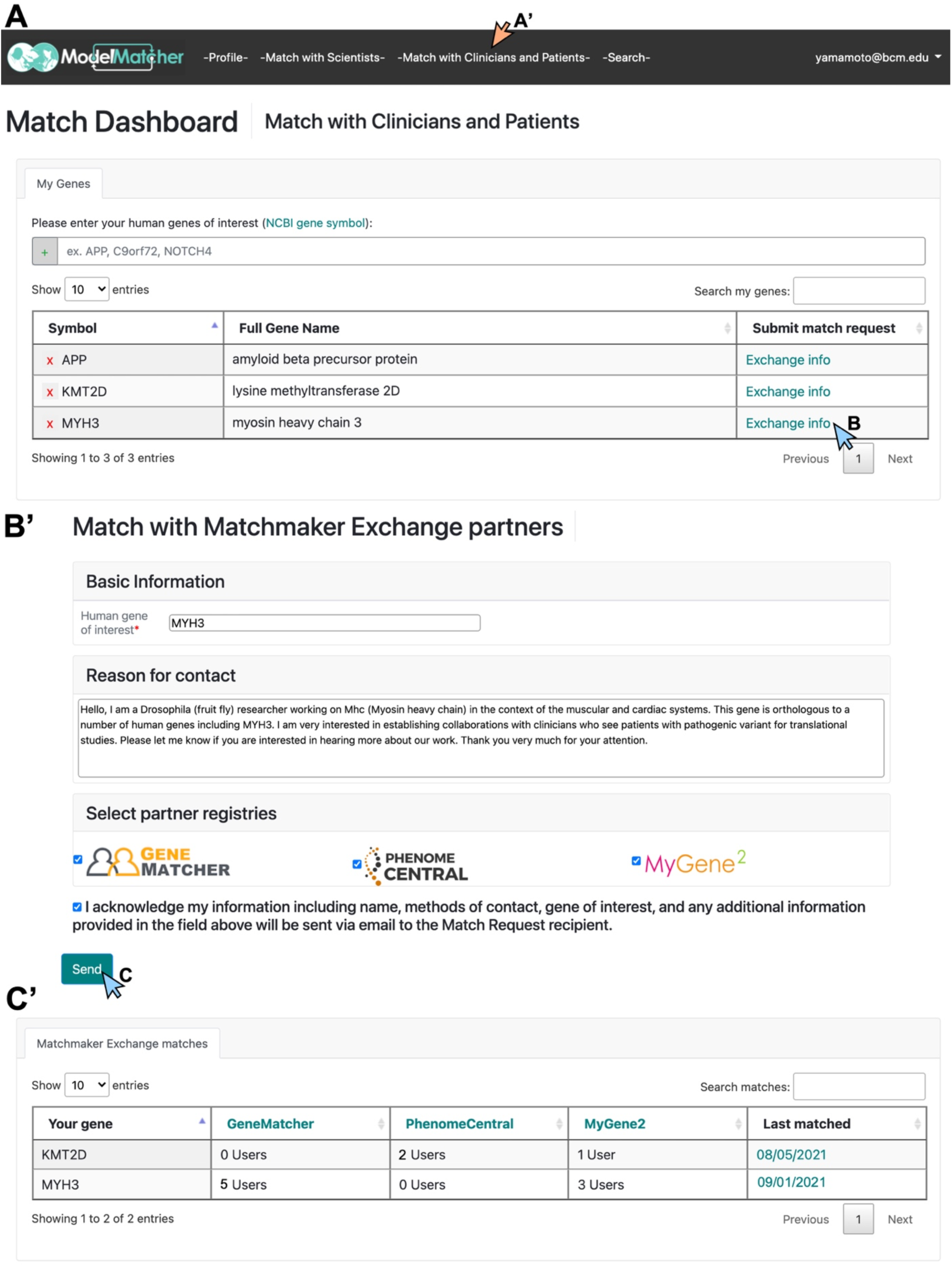
ModelMatcher users can actively identify potential clinical collaborators through Matchmaker Exchange. **A)** By logging in to their Match Dashboard account and clicking the -Match with Clinicians and Patients- button (arrow in **A’**), users will be taken to a tab to identify clinicians and patients who are registered in our partner registries in Matchmaker Exchange (currently GeneMatcher, MyGene2, and PhenomeCentral) who share interests in the same gene. The list of the user’s human genes of interest, which were previously entered into the Match Dashboard through the -Match with Scientists- tab, will be automatically loaded. Additional genes can also be added here. **B-B’)** By clicking the “Exchange Info” button next to each gene in **A)**, a Match Request Form to exchange information with clinical users of partner registries will appear. The User will be asked to briefly explain why they wish to connect with clinicians and patients who express interests in the specific gene, select the specific databases that they wish to exchange information with (default setting is to attempt to match with all partners), and consent that the information provided in this form and their user profile will be sent to the other party. **C)** By clicking the “Send” button, the Match Request will be sent to the corresponding partner registries and the user will receive an email regarding their matches. **C’)** The information regarding each Match Request sent by the user will also be archived in the “Clinician and Patient Matches” box at the bottom of the -Match with Clinicians and Patients- tab. Note that the Match results displayed here are synthesized data from test servers and do not reflect the actual matches with clinicians and patients who participate in Matchmaker Exchange.

Reciprocally, MME nodes can send match requests to ModelMatcher if their users wish to identify scientists with expertise in certain human genes or their ortholog candidates. ModelMatcher uses the information stored within the Scientist Registry to identify matches and provides information to both the requester and the matching scientist. When users from other registries send Match Requests through the MME, they are typically be given an option to opt-in to match with a ModelMatcher scientist to discuss collaboration opportunities. Using the aforementioned ortholog conversion standard, ModelMatcher converts human gene queries from MME into the corresponding model organism genes to identify matching scientists. Depending on the privacy level set for the specific gene, we send back the full user profile or an anonymized contact form (similar to a Match Request Form to an Anonymous Scientist) to the MME partner registries. Users in our Scientist Registry will also immediately receive an email if they appear in an external Match Request, providing them with the information on the originating registry, relevant case information, and contact information of the other party to help facilitate a decision on potential collaborations.

## Discussion

Implementation of next generation sequencing technologies into clinical care over the past decade has revolutionized rare disease diagnosis. However, as clinicians and researchers sequence more patients, they find more and more VUS in known disease-causing genes, or rare potentially damaging variants in GUS. These cases need functional assessment through experimental strategies to evaluate the pathogenicity of the disease-associated genetic variants. This necessitates the creation of collaborative research platforms that can connect patients, clinicians, and scientists. While multiple efforts to facilitate experimental studies of genetic variants identified in rare disease patients have been launched in different parts of the world with great success, many scientists with expertise in specific genes and biological processes who can make significant contributions do not have the opportunity to contribute. This often happens because the scientists are not part of the collaborative consortia or networks due to several barriers to entry. ModelMatcher is a network designed to place individual scientists at the center of our registry. We specifically designed this platform so clinicians, patients, and other stakeholders of rare and undiagnosed disease research can easily access scientists, who may reside in other countries, and initiate the collaborations needed to solve rare disease cases. Reciprocally, by connecting ModelMatcher to MME, we provide a way for scientists to reach out to clinicians and patients, which can help increase the clinical significance of their research projects. Through these efforts, we hope that more researchers working in pure-scientific laboratory settings will gain better access and stronger connections to the medical community. The integration of ModelMatcher to the larger RDMM International Network further increases the number of scientists who can participate and contribute to this ecosystem. This will have long-term beneficial effects by both bolstering rare disease research and advancing the fields of basic or fundamental biology.

There is well established data that shows how beneficial an accurate diagnosis is to the mental health of patients and family members, regardless of how it might change patient management (McConkie-Rosell et al., 2018). Additionally, there are cases in which identifying the affected gene has led to new treatment strategies being implemented. Over the years, many patient and patient support groups have also formed as people with a diversity of undiagnosed diseases receive genetic diagnoses and reach out to other families who received a similar diagnosis. Family conferences hosted by these groups allow for a more holistic view of symptoms across patients, which can be variably expressed and incompletely penetrant, to deepen our understanding of the newly documented condition. In addition, diagnosis opens up the opportunity for coalition building around a certain genetic condition, forming a community that can support one another and grow as a group as more patients become recognized with the same or related disorders. This, combined with group lobbying and fund-raising efforts, can facilitate resource allocation to support biomedical research on the genes and diseases of interest, increasing biological knowledge that can be used to design effective treatments and therapy. ModelMatcher can facilitate this effort by matching patients’, clinicians’, and scientists’ interests in the same or orthologous genes to make appropriate connections in order to boost the speed of translational and preclinical research.

Clinicians, patients, and support organizations are not the only groups to benefit from connecting with scientists. There are several stakeholders that are not often considered in the current collaborative matchmaking paradigm. Government agencies, non-profit organizations, biotech and pharmaceutical companies, and venture capital funds all desire to make informed decisions about who to contract for specific projects, and where to allocate their funds. A common strategy for these groups is to mine publications to find scientists and clinicians working on specific genes or pathways related to their project using online search tools such as PubMed (https://pubmed.ncbi.nlm.nih.gov/). This is suboptimal for two reasons: scientists who have worked on a gene in the past do not necessarily continue to work on it, and there could be scientists currently studying a gene who have not yet published their findings (**Table 1**). Having a central, up-to-date database like ModelMatcher will streamline collaborator identification and fund allocation to the appropriate scientists to accelerate specific projects.

While collaborations can bring tremendous benefits to biomedical research and clinical care, there are pitfalls that must be carefully avoided when working with others. Those unfamiliar with collaboration, especially across disciplines, should be mindful of expectations; both the expectations they have of others and the expectations others place on them. This is especially true for any users coming into ModelMatcher, as this service is designed solely to facilitate collaboration initiation, but does not provide guidance to maintain and expand collaborations once they are established. Collaborations between scientists and clinicians will not universally lead to diagnosis or treatment due to several factors. Even when they do, the timeline for these developments can be years long. All parties entering a collaboration should discuss realistic timelines, expectations, and what information will need to be shared prior to embarking on a project. All parties should be responsible for analyzing and vetting clinical and scientific information provided by other groups to ensure the project stands on solid foundations. Communication amongst members of the collaboration should be honest, open, and frequent. Considering that many ModelMatcher users are likely to be collaborating with people with different backgrounds and interests for the first time, it is crucial that everyone puts a significant amount of effort and energy into the project so the collaboration goes smoothly. Each participant must remember that a bad collaboration experience may not only ruin the current project but could also poison the collaborative atmosphere around them and make others hesitant to take part in future cross-disciplinary collaborations.

Finally in addition to its impact on the rare and undiagnosed research community, ModelMatcher can be used to facilitate research on common diseases. WES and WGS are being used to genotype patients with common diseases such as type 2 diabetes (Xue et al., 2018), autism spectrum disorders (Grove et al., 2019) and Alzheimer’s disease (Jansen et al., 2019). While these disorders tend to be more complicated than rare diseases because multiple genetic and environmental factors often contribute to disease risk and expressivity, functional studies of disease associated variants are often desired or required. Moreover, interpretation of somatic mutations identified in cancer patients can also be facilitated by functional studies performed by scientists, including model organism biologists who can perform gene and protein specific assays *in vivo* (Ablain et al., 2018; Bangi et al., 2019). Importantly, studies of rare diseases can provide important biological insights into related common diseases since some rare and common diseases converge on the same or similar pathogenic mechanisms. Finally, rare genetic disease patients often provide valuable knowledge about *in vivo* gene function in human, which cannot be obtained through model organism research or through *in vitro* or *ex vivo* studies performed in test tubes. The informatic resource available through ModelMatcher has numerous applications, which we hope will benefit the larger biomedical research community to improve healthcare and advance science for mankind.

## Acknowledgements

The ModelMatcher project is supported by the Jan and Dan Duncan Neurological Institute at Texas Children’s Hospital, a grant from the National Institutes of Health (NIH, U54NS093793) to M.F.W., H.J.B, Z.L. and S.Y., and support from the Hamill Foundation to L.L.. The Undiagnosed Diseases Network (UDN) is supported by the NIH Common Fund. H.J.B received support from the Howard Hughes Medical Institute. The Rare Diseases Models and Mechanisms (RDMM) Network is funded by the Canadian Institutes of Health Research (RCN-137793 and RCN-160422) with additional support from Genome Canada and Genome British Columbia to K.M.B. and P.H.. We thank the leadership and technology teams of Solve-RD, Australian Functional Genomics Network, GeneMatcher, MyGene2 and PhenomeCentral for data sharing and technical advice. We thank Dr. Huda Y. Zoghbi and the leadership teams of Alliance of Genome Resources (AGR), UDN Participant Engagement and Empowerment Resource (UDN PEER), and Matchmaker Exchange for valuable advice and useful discussions. We also acknowledge individual scientists for their active participation in ModelMatcher and the RDMM International Network, and clinicians, patients, and family members for their valuable contributions to research initiatives that are part of the Matchmaker Exchange. We thank Mr. David Deen for icon and logo design. The authors have no conflict of interest to declare.

## Methods

ModelMatcher is hosted on an AWS EC2 virtual machine running Ubuntu 18.04 and consists of four components: the landing/information page, the Scientist Registry, the Match Dashboard Registry, and MME API endpoints. The Scientist Registry operates a modified version of the RDMM server application (Boycott et al., 2020). The landing/information page, Match Dashboard Registry, and API endpoints operate as a single Node.js/Express application. Domain hosting and DNS services are provided by domain.com.

### Landing/Information Page

The landing page of ModelMatcher is an HTML-based web page that that is opened when a user visits our website using internet browsers (https://www.modelmatcher.net/). The page contains general information about the project (“About us” tab), a description of how the platform works (“Instructions” tab), answers to frequently asked questions (“FAQ” tab), contact form to reach the administrative team (“Contact” tab), a page with hyperlinks to our partner registries and related websites (“Links” tab), a link to page to register or log in as a Scientist or a Match Dashboard user (“Register/Login” tab), and a “Search” button to search information on the Scientist Register. The “Links” tab also contains a hyperlink to a webpage that allows users to download icons representing model organisms, organ systems and scientific categories that were developed for this project (https://www.modelmatcher.net/downloads.html). These icons can be freely used under Creative Commons license BY-NC 4.0. Hyperlinks to social media accounts are also provided, which provides tutorial videos (YouTube: https://www.youtube.com/channel/UC2o1FuMMZoa6oe_ekphd0sw and updated news (Twitter: https://twitter.com/modelmatchernet; Facebook: https://www.facebook.com/modelmatcher).

### Scientist Registry and the Search feature

The Scientist Registry of ModelMatcher is based on version 1.4.3 of the RDMM registry application hosted on GitHub (https://github.com/pavlidisLab/rgr). The program is primarily written in Java, with Spring and Thymeleaf providing the server functionality. The backend database service utilizes MySQL 8. Gene lists and metadata for each model organism were downloaded from the NCBI gene information database (https://www.ncbi.nlm.nih.gov/gene) and entrez gene IDs are used for internal identification purposes. Uberon is used for organ system metadata information (Mungall et al., 2012). Additional information can be found on the linked GitHub page shown above, or in the RDMM publication (Boycott et al., 2020).

We implemented the RDMM registry application for ModelMatcher with very limited changes. The only major modification was the implementation of a stricter ortholog candidate filtering criteria, necessitated by the lack of manual review in ModelMatcher compared to RDMM. We implemented a filter based on “DIOPT scores greater than 5” and “either best forward or reverse match” based on DIOPT version 8.0 (Hu et al., 2011). Some ortholog pairs with low or no DIOPT scores were manually entered. The latest ortholog candidate conversion table can be downloaded from our website (https://www.modelmatcher.net/assets/homologs.csv). All other modifications were minor, involving either text or visual styling changes.

The Search function of ModelMatcher is based on the RDMM’s initial implementation. Through the RDMM API, ModelMatcher users can discover scientists in partner registries that are part of the RDMM International Network (currently RDMM, Solve-RD, AFGN), and ModelMatcher scientists can also be discovered by users of our partner registries.

### Match Dashboard Registry and the Match Request Form

The backend of the Match Dashboard Registry uses MongoDB for user registration and authentication and accesses the Scientist Registry MySQL database for matching and API queries. We use the Node.js Passport package to handle user authentication and login and the Nodemailer package to provide email server capabilities.

Match Dashboard is a user-based system that requires login to ensure security and allow users to maintain a long-term profile complete with genes of interest. Unlike the Scientist Registry, genes of interest stored in the Match Dashboard Registry are not shared with other users, unless the user actively submits a match request to MME. To identify a potential scientist collaborator, users enter their human genes of interest in the -Match with Scientists-tab. ModelMatcher converts human genes of interest entered by the Match Dashboard users to appropriate ortholog candidate using the conversion table as described above and access the Scientist Registry SQL database to query for matches. These matches are presented to the user in a table format. The user can either manually send an email based on the contact information displayed, or use the provided Match Request Form to send an email through the backend of ModelMatcher. User profile state is maintained so that match requests sent through the backend have the most vital information (e.g., name, contact email) automatically populated. The account-based structure also allows future development of additional features such as sending notification emails to users when a new scientist has registered a gene of interest.

### Matchmaker Exchange API endpoints

The MME API endpoints use version 1.1 of the MME standard specification (https://github.com/ga4gh/mme-apis). For incoming queries, we use only the human gene symbol for matching purposes with our match strength score determined by DIOPT scores. We query the Scientist Registry for matches based on the previously described ortholog candidate conversion table. We return the corresponding information of our Scientist Registry users, and also provide a link to their profile information. If the gene of interest is set to the Restricted privacy category by the user, we provide more limited profile information (PI status, model system, model organism, organ system of interest) and a link to a contact form (similar to the Match Request Form) to email the Anonymous Scientist through the backend without needing to create a ModelMatcher account. Whenever we provide data to an external partner, we also notify our own users and provide them with the matching information we receive through the query. Match Dashboard users can also request matches to participants of partner MME registries (currently GeneMatcher, MyGene2 and Phenome Central) through the -Match with Clinicians and Patients- tab. When a match is initiated, their profiles and case information are shared through the API. We notify any matches based on a standardized email template. The match history is also stored in their user profile and displayed on the -Match with Clinicians and Patients- tab.

## Supplemental Materials

**Supplemental Table 1: List of websites and internet links cited**

**Supplemental Table 2: Consortia member list (Members of the Undiagnosed Diseases Network)**

